# Developmental changes in cerebral NAD and neuroenergetics of an antioxidant compromised mouse model of schizophrenia

**DOI:** 10.1101/2022.03.04.482878

**Authors:** Radek Skupienski, Pascal Steullet, Kim Q. Do, Lijing Xin

## Abstract

Defects in essential metabolic regulation for energy supply, increased oxidative stress promoting excitatory/inhibitory imbalance and phospholipid membrane dysfunction have been implicated in the pathophysiology of schizophrenia (SZ). The knowledge about the developmental trajectory of these key pathophysiological components and their interplay is important to develop new preventive and treatment strategies. However, this assertion is so far limited. To investigate the developmental regulations of these key components, we assessed, for the first time, *in vivo* redox state from the oxidized (NAD^+^) and reduced (NADH) form of Nicotinamide Adenine Dinucleotide (NAD), energy and membrane metabolites, neurotransmitters (*γ*-aminobutyrate and glutamate) by ^31^P and ^1^H MRS during the neurodevelopment of a SZ animal model with genetically compromised glutathione synthesis (*gclm*-KO mice). When compared to age-matched wild type (WT), an increase in NAD^+^/NADH redox ratio was found in *gclm*-KO mice until early adulthood, followed by a decrease in full adults as observed in patients. Especially, in early postnatal life (P20, corresponding to childhood), levels of several metabolites were altered in *gclm*-KO mice, including NAD^+^, ATP, phosphocreatine, intracellular pH, glutamine + glutamate and membrane phospholipids components, suggesting an interactive compensation for redox dysregulation between NAD, energy metabolism, and neurotransmission. The identified temporal neurometabolic regulations provide insights in preventive treatment targets for at-risk individuals, and other neurodevelopmental disorders involving oxidative stress and energetic dysfunction.

## Introduction

Schizophrenia (SZ) is a neurodevelopmental syndrome, affecting ~ 0.4% of the population, which is arising from both genetic and environmental factors ^1^. Although the disease is characterized by an increased dopamine release in the striatum (upregulation of the mesolimbic pathway) and a reduced release in the prefrontal cortex (downregulation of the mesocortical pathway) ^2^, other pathophysiological processes have been implicated: dysfunction during development in NMDAR-mediated signaling, neuroimmune regulation, or mitochondrial function appear to initiate “vicious circles” centered on redox dysregulation/oxidative stress, leading to anomalies of parvalbumin interneurons (PVI) and excitatory/inhibitory imbalance, at the basis of cognitive deficit ^3–5^. Additionally, defects in essential metabolic processes for energy supply ^6^ and membrane function ^7^ have also been implicated in SZ.

Glutathione (GSH), which is one of the main cellular antioxidants, has been reported to be reduced in the brain of some patients with schizophrenia ^8–10^. Proper redox homeostasis relies on the tight reciprocal interactions between the antioxidant systems and the energy metabolism which produces reactive oxygen species ^11^. The energy metabolism is dependent on many oxido-reduction enzymatic reactions involving co-agonist redox couples, such as NAD^+^ (oxidized) and NADH (reduced) forms of nicotinamide adenine dinucleotide (NAD). They act as cofactors in bioenergetic pathways and play a fundamental role in oxido-reduction reactions, such as glycolysis, oxidative phosphorylation, free radical detoxification, and superoxide production ^12^. The cellular oxidoreductive state can be estimated by the redox ratio (RR; NAD^+^/NADH). NAD^+^ is also involved as a co-substrate in many other biologically relevant processes, including calcium homeostasis, immunological functions, carcinogenesis, cell death, and gene expression ^13^. Moreover, the ratio NAD^+^/NADH has recently been shown to be decreased in the frontal lobe of patients with SZ ^14^, suggesting a redox imbalance in SZ. However, it is unclear how key pathophysiological components, including NAD redox status, cerebral energy metabolism, excitatory and inhibitory neurotransmission, membrane metabolism and their interplay, unfold from childhood to the onset of the disease. Such knowledge is important for developing new preventive and treatment strategies.

In this study, we aim to assess how a redox dysregulation caused by a deficit in GSH as seen in subsets of patients impacts RR, energy metabolism, excitatory and inhibitory metabolites along the postnatal development period. Previously, we demonstrated the feasibility of measuring RR, energy and membrane metabolites during mouse brain development using phosphorus magnetic resonance spectroscopy (^31^P-MRS) ^15^ at 14.1T, which is challenging because of the low sensitivity due to the small brain size. Thus, we pushed towards the characterization of developmental alterations in a mouse model related to SZ, which displays reduced GSH synthesis (*gclm*-KO) leading to several characteristic features of the disease, including impaired inhibitory GABAergic parvalbumin-positive interneurons (PVI) in numerous brain regions ^16^, neuroinflammation, compromised mitophagy ^17^, white matter and oligodendrocyte anomalies ^18,19^ together with altered behaviors ^20^. *Gclm*-KO mouse phenotype is characterized by a 60-70% decrease in GSH and a slightly lower weight as compared to wild-type control (WT). In addition, its blood glucose, as well as liver glycogen are lower, indicating a sustained metabolic activity where the sugar is directly processed ^21^. This mouse also depicts a redox imbalance leading to increased oxidative stress together with a greater vulnerability to external stresses ^22^.

Therefore, we focused on the measurement of frontal NAD^+^ and NADH content, energy metabolites, neurotransmitters (*γ*-aminobutyrate and glutamate) by ^31^P and ^1^H MRS during the development of *gclm*-KO and WT mice to investigate the developmental trajectory of these key components and to assess whether the mechanistic links between these processes along brain development may be relevant for the pathophysiology of SZ.

## Materials and Methods

### Study plan

To have an overview of the critical time point in neurodevelopment implicated in schizophrenia, a cohort of mice was scanned at different ages corresponding to different developmental stages (Fig. 1a): Postnatal day 20 (P20, the end of the suckling period); P40 (the puberty period); P90 (the end of adolescence or early adulthood usually corresponding to the first appearance of psychotic episode in SZ) and finally P250 (mature adult)^23–25^.

**Fig. 1:**
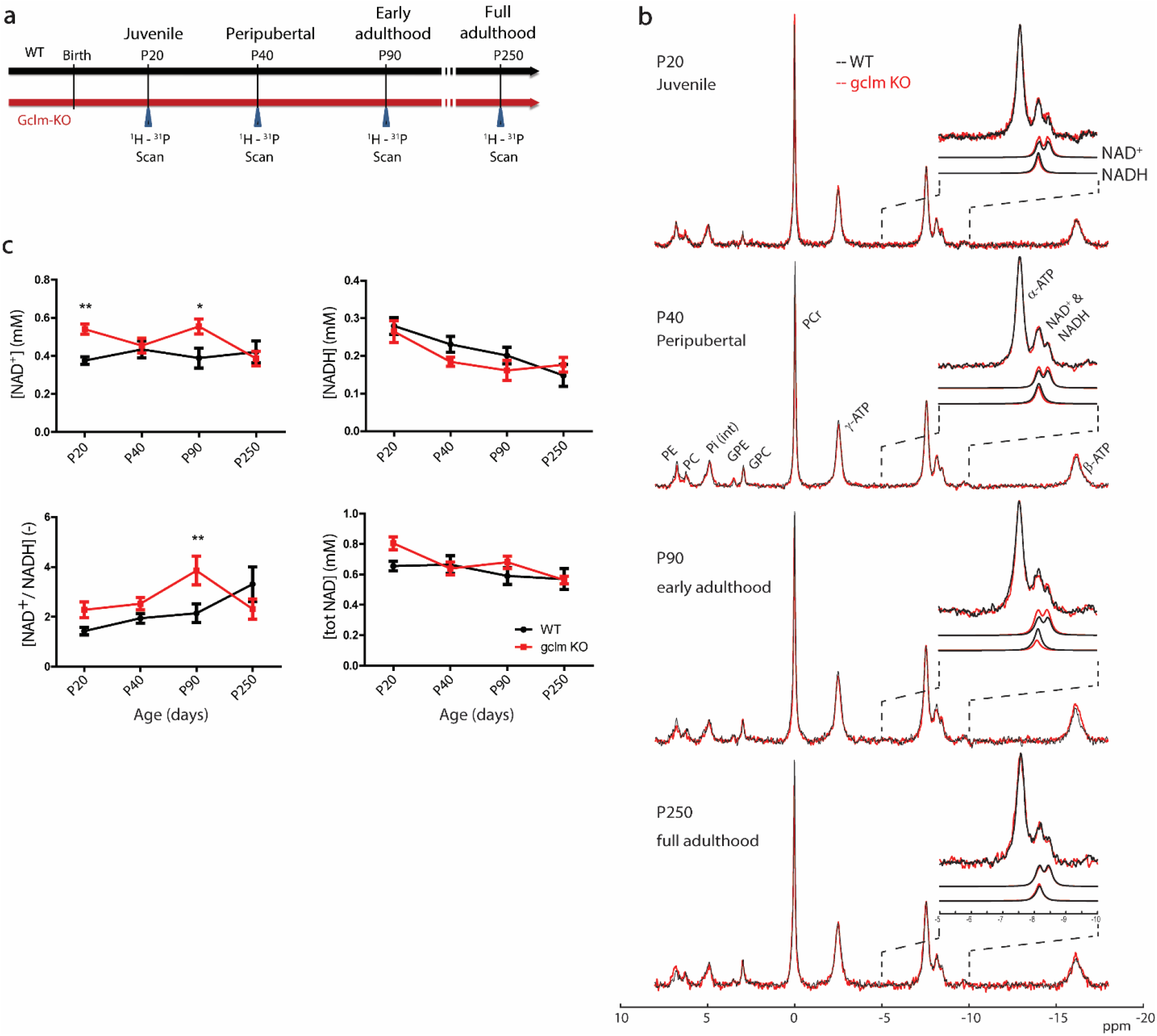
Developmental changes of cerebral NAD content and NAD^+^/NADH ratio in gclm-KO and WT mice. Changes of NAD content and redox ratio in *gclm*-KO and WT mice during brain development. **a** Study scheme presenting the different time-points and their corresponding developmental stages at which mice were scanned. **b** Summed ^31^P-MRS spectra (see the zoom on α-ATP and NAD spectral region) where the developmental changes of NAD^+^ and NADH can be visually observed along age (from P20 to P250) in WT and *gclm*-KO mice. **c** NAD^+^, NADH, RR and total NAD levels from P20 to P250 in WT and *gclm*-KO mouse brain. Significant increase of NAD^+^ at P20 and P90, and NAD^+^/NADH at P90 as compared to age-matched WT. At P250, a decrease of RR is observed in the *gclm*-KO as compared to P90, while in WT a stable NAD^+^ and decrease in NADH lead to an increase in RR.

### Animal model

*Gclm*-KO mouse model was generated and kindly provided by TP. Dalton (Cincinnati University, Ohio). The animals were frequently backcrossed with WT C57BL/6J from Charles River Laboratories to avoid genetic drift. *Gclm*-KO and WT mice born in the local animal facility were housed in ventilated cages on a 12-hour light-dark cycle at room temperature of 20-22 °C with 50-60% humidity. After weaning at P21, a minimum of 2 and a maximum of 5 animals were kept per cage. Tap water and regular chow were provided *ad libitum*. The experiments were conducted with a cohort of males/females, aged from 20 to 250 days with a bodyweight of 7-40 g. Respective animal numbers, age, genotype, males/females were as follows: P20, WT (5m/6f) *gclm*-KO (6m/6f); P40, WT (4m/5f) *gclm*-KO (5m/3f); P90, WT (4m/5f) *gclm*-KO (4m/5f); P250, WT (4m/1f) *gclm*-KO (3m/1f).

Animals were anesthetized with isoflurane (0.9 −1.2%) in a mixture of air and O_2_ (50/50%). The head of the animal was then fixed in a mouse holder with a bite bar and two ear inserts (RAPID Biomedical GmbH, Rimpar, Germany). The body temperature was kept at 37.0 ± 0.5 *°*C by tubing with circulating warm water controlled with a rectal probe. Spontaneous breathing was maintained at 90 ± 20 rpm by adjusting the isoflurane concentration. The respiration rate and body temperature were monitored by a small animal monitor (SA Instruments Inc., Stony Brook, NY, USA). All animal procedures were performed according to federal guidelines and were approved by the Swiss cantonal veterinarian authorities.

### In vivo ^1^H and ^31^P MR Spectroscopy

All MR experiments were performed on a 14.1 T small animal scanner with a 26 cm horizontal bore (Magnex Scientific, Abingdon, United Kingdom), equipped with a 12 cm internal diameter gradient coil insert (400 mT/m, 120 μs) and a DirectDrive console interface (Agilent Technologies, Palo Alto, CA, USA). Radiofrequency transmission/reception was achieved using a homebuilt three-channel surface coil, namely two single-turn loops (10 mm diameter) 90° geometrically decoupled, quadrature ^1^H coil with one loop linearly polarized ^31^P coil (10 mm diameter). Fast spin-echo multi-slice images were first acquired for voxel positioning using the following parameters: repetition time of 3.3 s, echo time of 43.24 ms, echo train length of 8, echo spacing of 10.81 ms, field of view of 20 x 20 mm, matrix size 128 x 128, slice thickness 0.4 mm, 35 slices and 2 averages. Local shimming in the volume of interest (VOI) was achieved using 1^st^- and 2^nd^-order shims with FAST(EST)MAP ^26^.

Water suppressed ^1^H MR spectra were acquired in the anterior cingulate cortex (ACC) from a volume of 5.76 μL (0.9 x 4 x 1.6 mm) using a SPECIAL sequence with an echo time of 2.75 ms, repetition time of 4 s, and 240 scans (30 x 8 blocks). The transmitter frequency was set on the water resonance for the acquisition of water spectra (8 scans) for metabolites quantification and eddy current correction. For normalization of the ^1^H MRS, the brain water content was measured at P20, P40, and P90 using the weight difference between freshly removed and fast dissected brain structures, and its residue after lyophilization (Fig. S1). The water content at a later stage (P250) was assumed to be stable and the same as that at P90.

^31^P MR spectra were acquired in the fronto-dorsal part of the brain using a 3D-ISIS localization in combination with a pulse-acquire sequence composed by an adiabatic half passage pulse (500 μs pulse width) with the transmitter offset was set on NAD^+^ (−8.3 ppm). The following measurement parameters were used: voxel size 90 μL (2.5 x 6 x 6 mm^3^) at P20 and P40, 122.5 μL (2.5 x 7 x 7 mm^3^) at P90 and P250, TR = 5 s, 1600 averages (100 blocks of 16 averages), 12 kHz spectral width and 4096 complex points.

### Spectral quantification

Frequency drift and phase variation were corrected prior to the summation of all block spectra. The ^1^H and ^31^P metabolite concentrations were determined with LCModel (Stephen Provencher Inc., Oakville, Ontario, Canada). Cramér-Rao lower bound (CRLB) was used as an exclusion criterion for all ^1^H and ^31^P data and the cutoff was set at a maximum of 30 %.

^1^H basis set containing a measured macromolecule spectrum and simulated metabolite spectra was used together with the unsuppressed water signal measured from the same VOI as an internal reference for the absolute quantification of metabolites. The following 21 metabolites were included in the analysis: acetate (Ace), alanine (Ala), ascorbate (Asc), aspartate (Asp), Creatine (Cr), *γ*-aminobutyrate (GABA), glucose (Glc), glutamine (Gln), glutamate (Glu), glycine (Gly), glycerophosphocholine (GPC), glutathione (GSH), myoinositol (Ins), lactate (Lac), N-acetyl-aspartate (NAA), N-acetyl-aspartylglutamate (NAAG), phosphorylcholine (PCho), phosphocreatine (PCr), phosphoethanolamine (PE), scylloinositol (Scyllo), taurine (Tau).

^31^P metabolite concentrations were normalized using PCr levels obtained from the ^1^H experiments. Apodization with a 10 Hz exponential function was applied to all spectra prior to spectral quantification. The ^31^P metabolites quantification was effectuated with a basis-set prepared with simulated ^31^P spectra including PCr, *α*-ATP, *β*-ATP, *γ*-ATP, Pi^int^ (intracellular inorganic phosphate), Pi^ext^ (extracellular inorganic phosphate), PE (phophothanolamine), PC (phosphocholine), GPC (glycerophosphocholine), GPE (glycerophosphoethanolamine), MP (membrane phospholipid), NADH, NAD^+^, with respective linewidths ^15^. Uridine diphosphate glucose (UDPG) is a phosphorylated sugar intermediate in biochemical reactions, and it contains resonances overlapping with NADH and NAD^+^. Due to the low concentration of UDPG and low sensitivity in mouse brain, additional analysis was performed with summed age pooled spectra with the inclusion of UDPG in the basis-set to quantify its level and to evaluate its effect on NAD quantification results. Physiological parameters (pH^int^ and free [Mg^2+^]) were calculated from chemical shift differences between ^31^P metabolites (details of the calculation can be find herein ^15^).

### Statistical analysis

All statistical analyses were performed in Graphpad Prism 5 (GraphPad software, inc.), Matlab (R2017a) or R (R Core Team, 2019). All variables were tested by two-way ANOVA using age and genotype as a fixed factor for the comparison between groups (*gclm*-KO vs. WT) or were tested by one-way ANOVA using age as a fixed factor for differences within the same genotype. In case of significant difference between groups, the effect of age was *post hoc* investigated between groups using the Bonferroni correction for multiple comparisons. In order to evaluate the presence of regular increase or decrease with age that would not be detected by ANOVA, linear regressions along age were performed. Correlations between all metabolites were effectuated using two-tailed Pearson correlation with 95% of the confidence interval and reported using the correlation coefficients R^2^. The results are presented as the mean and standard error of the mean unless otherwise stated. When exact p-values are not provided, significant differences (*) were considered for P < 0.05, (**) for P < 0.01, (***) for P < 0.001 and (****) for P < 0.0001.

## Results

### NAD content during neurodevelopment

NAD content in mouse fronto-dorsal brain was monitored *in vivo* by ^31^P MRS at four time points corresponding to different postnatal developmental periods (Fig. 1, Table 1). The summed ^31^P MR spectra, from P20 to P250, illustrate the differences between WT and *gclm*-KO (Fig. 1b) together with the excellent sensitivity and spectral quality at 14.1T. A zoom on the NAD spectral region allows to visually recognize the temporal changes of NAD^+^ and NADH in both genotypes.

**Table 1.** Concentration and CRLBs of NAD^+^, NADH, total NAD, and RR at P20, P40, P90 and P250, respectively.

| AGE<br>[DAYS] | GENOTYPE | NAD<sup>+</sup><br>[mM] | CRLB<br>[%] | NADH<br>[mM] | CRLB<br>[%] | RR<br>[-] | TOTAL NAD<br>[mM] |
| --- | --- | --- | --- | --- | --- | --- | --- |
| **P20** | WT | 0.38 ± 0.06 | 10 ± 1 | 0.28 ± 0.07 | 11 ± 2 | 1.43 ± 0.47 | 0.66 ± 0.10 |
|  | *Gclm*-KO | 0.54 ± 0.08 | 9 ± 2 | 0.27 ± 0.09 | 14 ± 3 | 2.28 ± 0.99 | 0.81 ± 0.13 |
| **P40** | WT | 0.43 ± 0.13 | 9 ± 2 | 0.23 ± 0.06 | 12 ± 3 | 1.94 ± 0.56 | 0.67 ± 0.18 |
|  | *Gclm*-KO | 0.46 ± 0.11 | 8 ± 1 | 0.18 ± 0.03 | 13 ± 2 | 2.53 ± 0.70 | 0.64 ± 0.12 |
| **P90** | WT | 0.39 ± 0.15 | 10 ± 3 | 0.20 ± 0.06 | 14 ± 3 | 2.15 ± 1.06 | 0.59 ± 0.16 |
|  | *Gclm*-KO | 0.56 ± 0.12 | 8 ± 2 | 0.16 ± 0.07 | 19 ± 4 | 3.85 ± 1.53 | 0.68 ± 0.12 |
| **P250** | WT | 0.42 ± 0.13 | 9 ± 1 | 0.15 ± 0.06 | 19 ± 5 | 3.31 ± 1.55 | 0.57 ± 0.15 |
|  | *Gclm*-KO | 0.39 ± 0.08 | 11 ± 1 | 0.18 ± 0.04 | 17 ± 3 | 2.31 ± 0.80 | 0.56 ± 0.05 |

^31^P MR spectra of individual animal demonstrated good spectral resolution (linewidth of PCr = 13-15 Hz) and good sensitivity (SNR = 30-50), which ensured the reliable quantification of the NAD signals with a mean CRLB of 9% for NAD^+^ and 15% for NADH. A significant difference between genotypes (two-way ANOVA with genotype and age as co-factors, P = 0.009) was observed for NAD^+^. However, the age did not have a significant influence on NAD^+^, while an interaction between age and genotype (P = 0.044) was found. The increase of NAD^+^ was found in *gclm*-KO as compared to WT at P20 (P < 0.01) and at P90 (P < 0.05) (Fig. 1c).

For NADH, no significant difference was observed between genotypes, but there was a strong age effect (P = 0.0002, Fig. 1c). Furthermore, a one-way ANOVA test within each genotype depicted a significant difference between ages (WT, P = 0.007; *gclm*-KO, P = 0.019). To characterize whether NADH linearly increases or decreases with age, a post test for linear trend was performed, which confirmed a significant decrease with age for both genotypes (WT, P =0.0008, R^2^ = 0.332, slope = −0.021; *gclm*-KO, P = 0.032, R^2^ = 0.140, slope = −0.014). *Post hoc* test showed a decrease between P20 and P250 in WT (P < 0.01) and between P20 and P90 (P < 0.05) in *gclm*-KO.

RR was shown to be affected by genotype (two-way ANOVA P = 0.044) and age (P = 0.006). An interaction between age and genotype was also found (P = 0.019). Elevated RR was found in *gclm*-KO compared to WT at P90 (P < 0.01). Further analysis using one-way ANOVA test on each genotype depicted a significant difference between age groups (WT, P = 0.006; *gclm*-KO, P = 0.030), and a post-test for linear trend showed a significant increase with age only in WT (P = 0.0006, R^2^ = 0.341, slope = 0.292). *Post hoc* test showed a RR increase between P20 and P250 (P < 0.01) in WT and between P20 and P90 (P < 0.05) in *gclm*-KO. Beyond P90, RR started to decrease in *gclm*-KO. Thus, to further characterize this observation, a segmented regression was performed (Fig. 2a). The segmented regression analysis (R Core Team, 2019), (corrected for repeated measures, age, and sex) depicted a switch at P90 in *gclm*-KO, with a positive slope from childhood to early adulthood and a negative slope from early adulthood to full adult (difference between both slopes, P = 0.005, Fig. 2b). In WT, no differences were found between these two slopes.

**Fig. 2:**
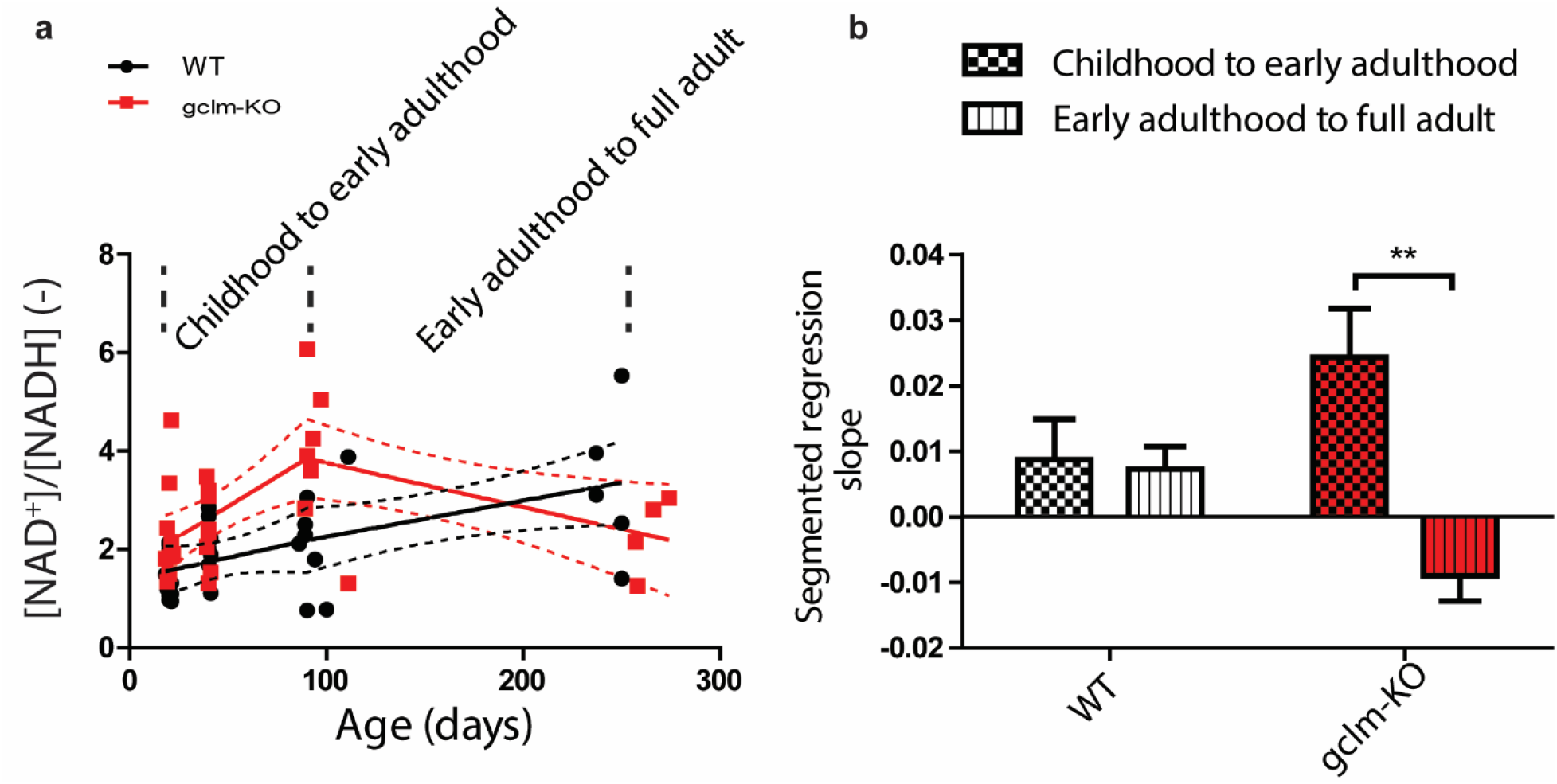
Switch of NAD^+^/NADH regulation in gclm-KO mice. Figure **a** show a stable increase of RR in WT, and an increase of RR until early adulthood followed by a decrease during adulthood in *gclm*-KO. **b** Slopes of the segmented regression clearly depict the difference between WT and *gclm*-KO mice (**p = 0.005), where a switch in RR occurs after early adulthood (P90) in *gclm*-KO, but not in WT.

Total NAD content did not depict any significant differences between genotypes but an age effect was observed (P = 0.020), which was driven by the *gclm*-KO group. Post-tests within individual genotype showed that total NAD remained unchanged in WT. However, a strong age difference (P = 0.006) was seen for *gclm*-KO. *Post hoc* test showed significant age-related decrease from P20 to P40 (P < 0.05) and from P20 to P250 (P < 0.05) together with a significant linear trend (P = 0.004, R^2^ = 0.232, slope = −0.034) decreasing along with age.

### Energy metabolites (ATP, PCr and Pi)

Genotype (P = 0.030), age (P < 0.0001) and the interaction between them (P = 0.010) had an influence on ATP levels. Significantly higher ATP level was observed in *gclm*-KO at P20 (P < 0.05) and at P90 (P < 0.05) in comparison to WT (Fig. 3). In WT, the ANOVA did not depict significant differences between age groups for ATP, but a slight age-related decrease was proven by the linear trend (P = 0.018, R^2^ = 0.172, slope = −0.092) along with age. In *gclm*-KO, ATP was significantly different between age groups (P < 0.0001). Post-test revealed a decrease from P20 to P40 (P < 0.0001), P20 to P90 (P < 0.01) and P20 to P250 (P < 0.01). A decreased linear trend with age was also observed (P = 0.003, R^2^ = 0.163, slope = −0.108).

**Fig. 3:**
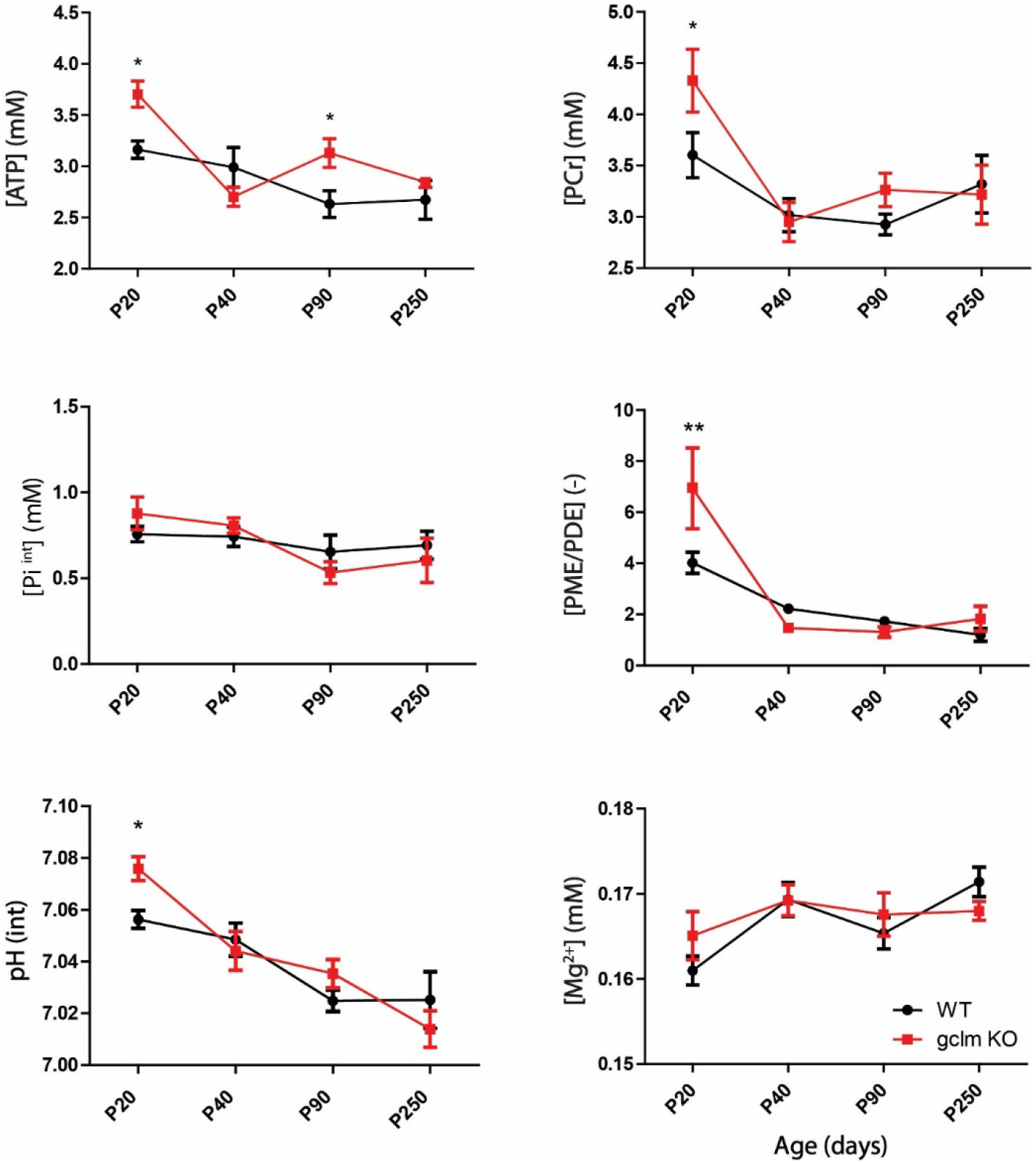
Brain developmental changes in ATP, phosphocreatine (PCr), Intracellular phosphate (Pi^int^), pH, [Mg^2+^] and phosphoester ratio PME/PDE in WT and gclm-KO mice. Changes of [ATP], [PCr], [Pi ^int^], pH, [Mg^2+^] and PME/PDE from postnatal day P20 to P250 in the brain of WT and *gclm*-KO mice. *Gclm*-KO mice demonstrated significantly increases in ATP, PCr, PME/PDE and pH levels at P20 relative to WT mice. All values are shown as mean ± SEM and significant differences are derived from the *post hoc* Bonferroni correction test for multiple comparisons. *P < 0.05, **P < 0.01.

A strong age effect was observed for PCr (two-way ANOVA, P < 0.0001). Although genotype difference was not detected by the ANOVA, the post-test revealed significantly higher PCr levels at P20 (P < 0.05) in *gclm*-KO in comparison to WT. In WT, no significant differences were observed with age. However, the age had a significant influence on PCr levels (P = 0.001) in *gclm*-KO. Thus, post-test highlighted a PCr decrease between P20 and P40 (P < 0.01), and between P20 and P90 (P < 0.05). A significant decreased linear trend was also observed (P = 0.020, R^2^ = 0.137, slope = −0.151). By contrast, Pi^int^ was rather stable across genotypes and age.

### Excitatory and inhibitory (E/I) metabolites (Gln, Glu and GABA)

No genotype difference was detected for Gln and Glu, but an age effect was found for Glu (P = 0.030). However for both Gln and Glu, an interaction between age and genotype was observed (P = 0.028 and P = 0.029, respectively). Both metabolites demonstrated a trend towards higher levels at P20 and lower levels at P40 in *gclm*-KO as compared to WT (Fig. 4). Furthermore, the analysis of their sum (Gln + Glu) presented neither genotype nor age differences, but a strong interaction (P = 0.0015). *Post hoc* analysis showed significantly more elevated Gln + Glu at P20 (P < 0.05), and lower Gln +Glu at P40 (P < 0.05) in *gclm*-KO in comparison to age-matched WT (Fig. 4). In WT, significant changes were observed with age (P < 0.0008), with post-hoc tests revealing a decrease of Glu + Gln between P20 and P40 (P < 0.001), and between P20 and P90 (P < 0.05). In contrast, the age did not influence Gln + Glu in *gclm*-KO mice. Finally, the concentrations of GABA did not differ between genotype and along with age.

**Fig. 4:**
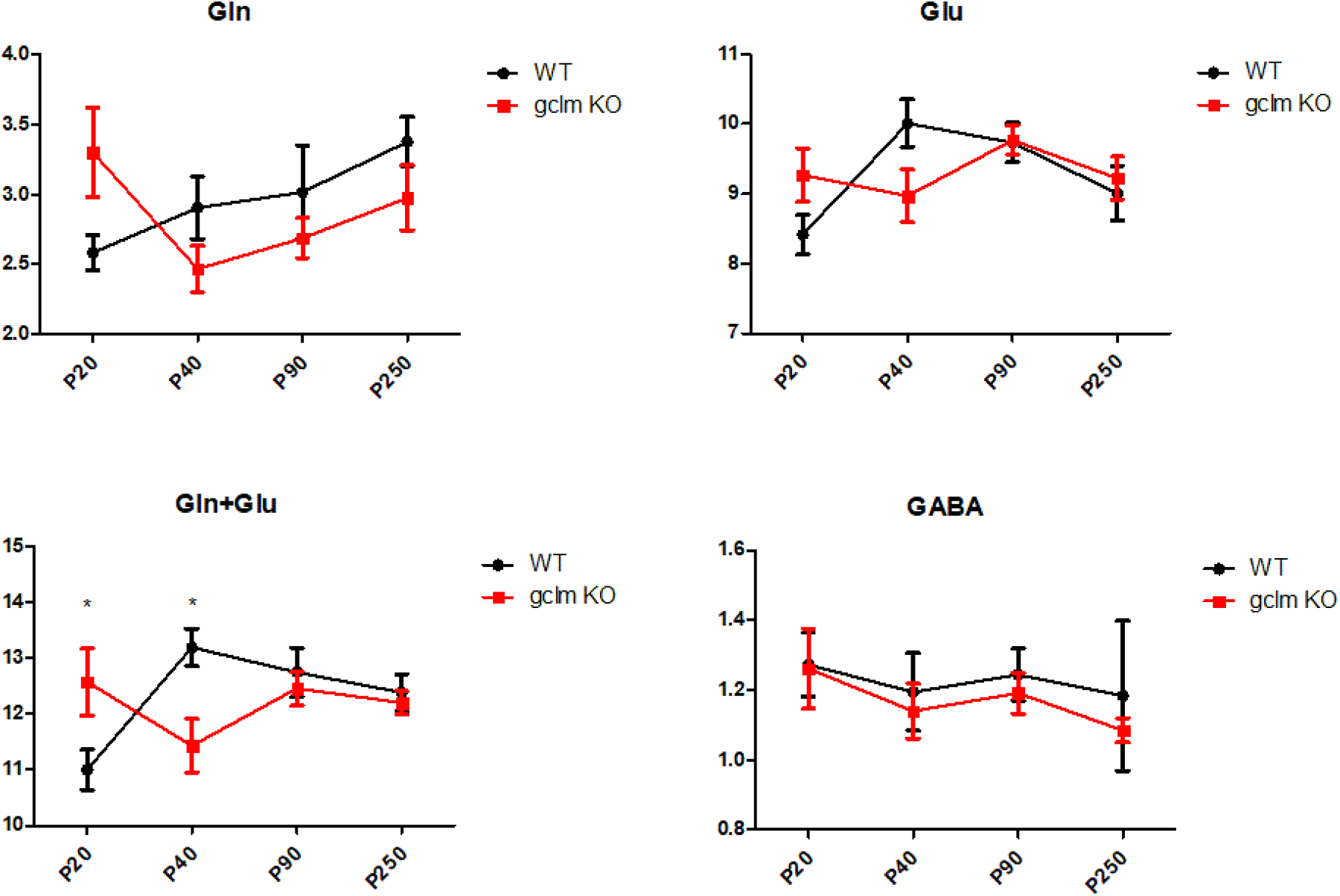
Excitatory and inhibitory (E/I) neurotransmitter concentrations (mM) in gclm-KO and WT mouse brain during development. Concentrations of E/I neurotransmission metabolites including Glu, Gln, Glu+Gln and GABA in *gclm*-KO and WT mouse brain during brain development. Strong trends of increases in Gln and Glu in *gclm*-KO at P20 lead to a significant increase of Glx (Gln + Glu). At P40, a significant reduction of Glx was observed in *gclm*-KO in comparison to WT.

### Association between NAD content, neurotransmission and energy metabolism

To investigate the interplay between NAD redox regulation, energy metabolism, and E/I neurotransmission, the correlations between the different components were analyzed at P20 and P90 which corresponded, in our study, to pivotal points of brain development (Fig. 5). P20 corresponds to the end of the suckling period after which a drastic change in alimentation happens together with the weaning. P90 matches with early adulthood in human, when the first psychotic episode occurs, and thus seems to represent a crucial transition phase from prodromal alterations to acute psychosis. At P20, NAD^+^ positively correlated with GABA (P = 0.009, R^2^ = 0.704), Gln (P = 0.016, R^2^ = 0.650) and PCr (P = 0.005, R^2^ = 0.755) in *gclm*-KO mice, but not in WT mice. NADH negatively correlated with Pi (P = 0.001, R^2^ = 0.739) only in WT. At P90, NADH positively correlated with ATP (P = 0.0006, R^2^ = 0.923) only in *gclm*-KO mice.

**Fig. 5:**
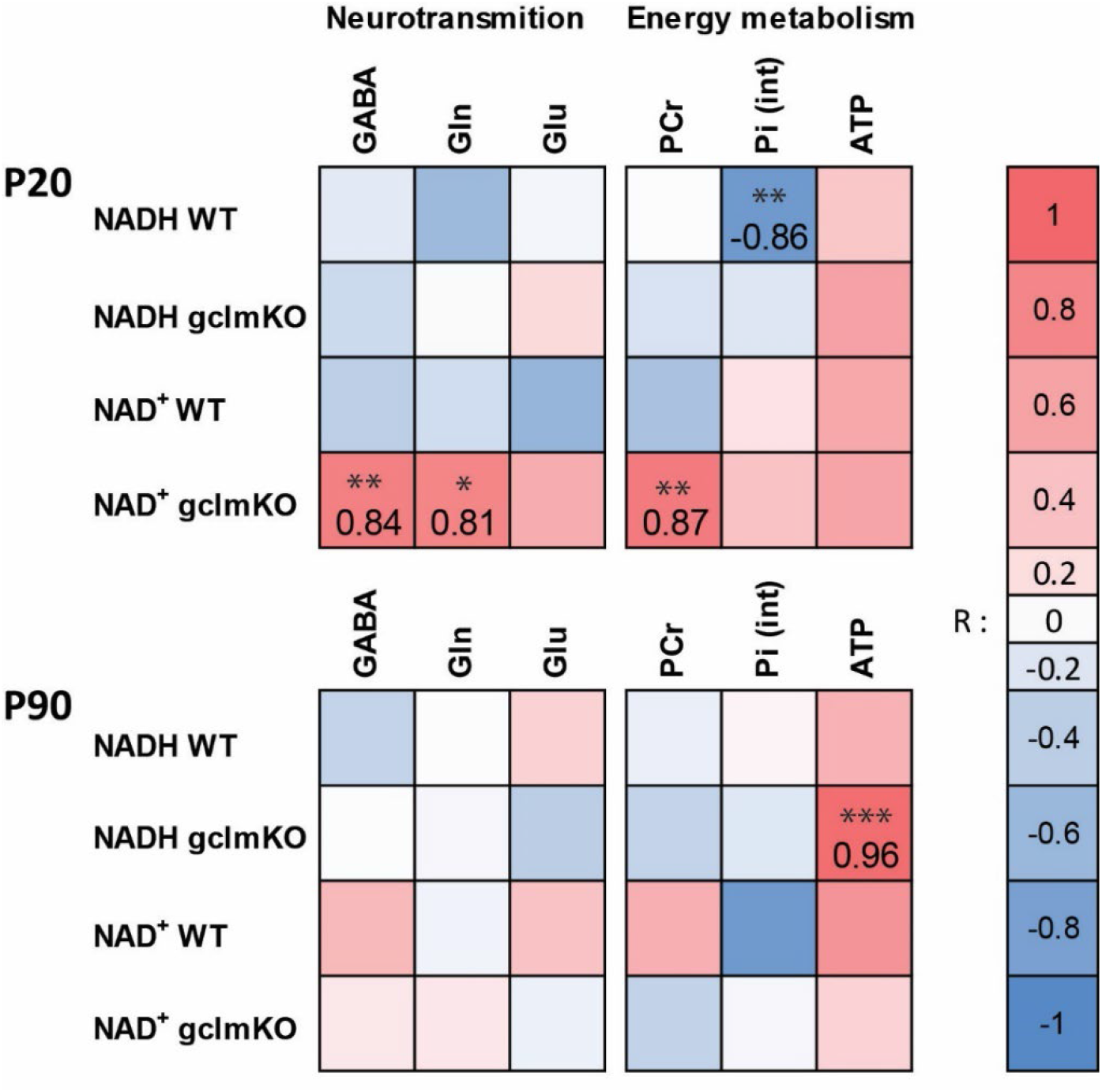
Associations between NAD content, E/I neurotransmission and energy metabolism. Correlation matrix depicts in *gclm*-KO and WT mouse the links between NAD content (NAD^+^ and NADH), neurotransmission and energy metabolism at P20 and P90. At early developmental period (P20), NAD^+^ is strongly associated with GABA, Gln and PCr in *gclm*-KO animals, while NADH is exclusively associated with inorganic phosphate in WT mice. At early adulthood (P90), *Gclm*-KO mice showed a strong positive association between their ATP and NADH levels. Heatmaps show the correlation coefficients. *P < 0.05, **P < 0.01, ***P < 0.001.

### Membrane metabolites

Phosphomonoesters (PME) (i.e. phosphoetanolamine and phosphocholine) that reflect membrane synthesis, and phosphodiesters (PDE) (i.e. glycero-phosphoetanolamine and glycero-phosphocholine) that reflect membrane degradation, were measured by ^31^P MRS (Fig. 3). Thus, the ratio (PME/PDE) represents the balance between membrane synthesis and degradation. No difference between genotype was detected in the PME/PDE ratio by two-way ANOVA, but a strong age effect was observed (P < 0.0001). Significant age differences were observed in WT (P < 0.0001) and in *gclm*-KO (P = 0.0009). Post-tests revealed a decrease in ratio from P20 to P40 (P < 0.001), P20 to P90 (P < 0.0001) and P20 to P250 (P < 0.0001) in WT, and from P20 to P40 (P < 0.01) and P20 to P90 (P < 0.01) in *gclm*-KO. A significant decreasing linear trend with age was also observed in both groups (WT, P < 0.0001, R^2^ = 0.463, slope = −0.439; *gclm*-KO, P = 0.012, R^2^ = 0.152, slope = −0.642). Post-hoc analyses revealed, however, a significantly higher ratio in *gclm*-KO at P20 (P < 0.05) in comparison to age-matched WT (Fig. 3).

### UDPG

UDPG is a phosphorylated nucleotide sugar intermediate of cell polysaccharide synthesis. By incorporating UDPG in the fitting prior knowledge, a significantly lower UDPG level (p < 0.05) was observed at P250 in *gclm*-KO mice (0.068 ± 0.031 mM), as compared to age-matched WT group (0.194 ± 0.059 mM). Due to the low level of UDPG, we further validated this result by summing all individual animal spectra at each age to improve the SNR. In line with results of the individual animal analysis, a three-fold decrease of UDPG was observed at P250 in *gclm*-KO relative to WT animals (Fig. 6, Table 1S).

**Fig. 6:**
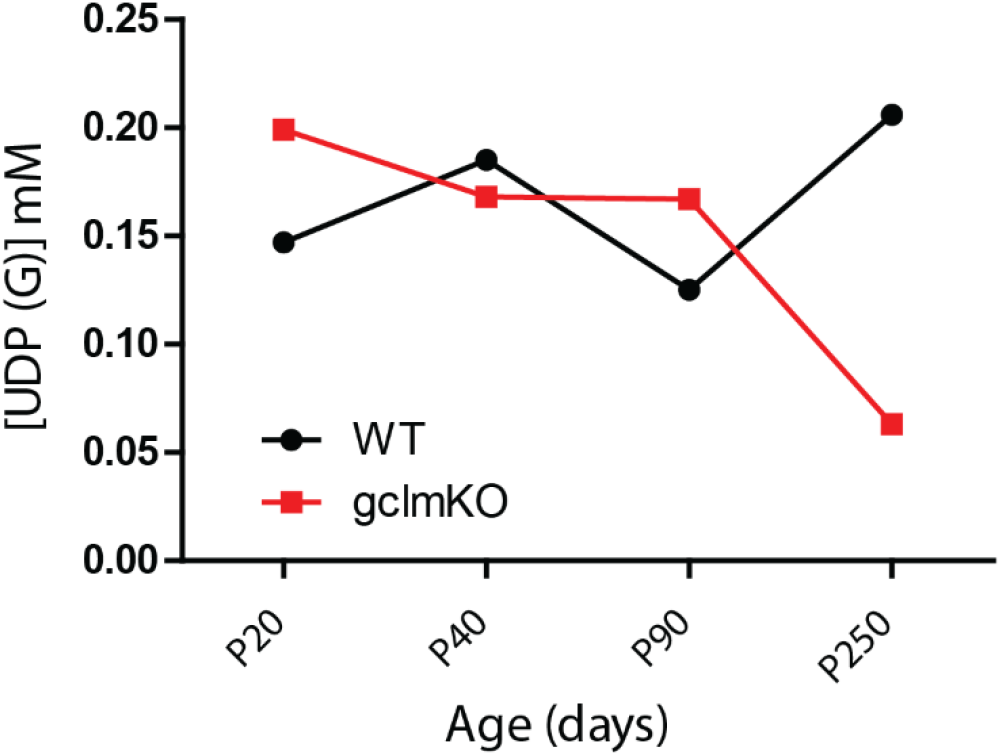
Levels of UDP(G) in WT and gclm-KO mice. Levels of UDP(G) in WT and *gclm*-KO mouse obtained from the pooled summed spectra, respectively at P20, P40, P90 and P250. Stable level along development was observed in WT mice, while a decrease after P90 occurred in *gclm*-KO mice.

### pH and [Mg^2+^]

pH and magnesium concentration [Mg^2+^] were also assessible by ^31^P MRS to monitor brain physiology. No difference between genotypes was detected by two-way ANOVA, but a strong age effect was observed (P < 0.0001). However, a post-hoc analysis highlighted a significantly higher pH^int^ (Fig. 3) in *gclm*-KO at P20 (P < 0.05) in comparison with age-matched WT. A separated one-way ANOVA analysis depicted age changes within each group (WT; P < 0.0009) and (*gclm*-KO; P < 0.0001). A significant decreasing linear trend with age was also observed in both groups (WT, P = 0.0003, R^2^ = 0.355, slope = −0.006; *gclm*-KO, P < 0.0001, R^2^ = 0.484, slope = −0.010). For [Mg^2+^], no difference was found between WT and gclm-KO, but an age effect was observed (P < 0.018). A significant increased linear trend with age was also observed in WT (P = 0.006, R^2^ = 0.195, slope = 0.001), but not in *gclm*-KO.

### Cerebral cortex water content

Cortical water content was measured in both WT and *gclm*-KO, at the different time points of interest (P20, P40 and P90) (Fig. 1S). A strong age effect was observed (two-way ANOVA, P < 0.0001) with a significant decrease between P20 and P40 for both genotypes (P < 0.0001), and between P40 and P90 for WT (P < 0.01). *Gclm*-KO displayed reduced water content at P40 (P < 0.05) in comparison to age-matched WT.

## Discussion and conclusion

With the sensitivity and spectral resolution enhancement at the ultra-high magnetic field (14.1T), using both *in vivo* ^31^P and ^1^H-MRS, we have identified, for the first time, the effects of a redox dysregulation due to genetically lower GSH levels on postnatal development of the NAD redox state, energy homeostasis, membrane metabolism, levels of excitatory and inhibitory neurotransmitters and their interactions in the prefrontal brain region. In *gclm*-KO mice, most alterations were observed during early life (at P20). This includes up-regulation of NAD^+^ resulting in higher RR, increases in Gln + Glu and membrane synthesis rate (increase in PME/PDE), and higher levels of the key energy metabolites PCr and ATP. In addition, NAD^+^ was positively correlated with GABA and Gln in *gclm*-KO at P20 but not in WT mice. At the end of adolescence/early adulthood (P90), the brain of *gclm*-KO mice also displayed concomitantly elevated levels of NAD^+^ and ATP as compared to age-matched WT mice. Noteworthy, a temporal evolution of NAD redox ratio during neurodevelopment was observed for *gclm*-KO mice. Thus, RR increased with age (from P20 to P90) with the highest value at P90, and then sharply dropped in older full adulthood *gclm*-KO, suggesting a shift from oxidative stress to reductive stress at adulthood. Such decline of NAD^+^/NADH in adult *gclm*-KO mice was similar to the one displayed by SZ patients ^14^, implying that a compensatory mechanism associated with the increase of NAD^+^ started to fail during adulthood as a consequence of a long-lasting, exacerbated oxidative stress.

### Developmental adaptation of NAD^+^ under oxidative stress

The *~* 70% decrease of brain endogenous antioxidant GSH in *gclm*-KO mice during the brain development induces oxidative stress in the ACC ^22^ and other sub-cortical structures ^16^. In the ACC of *gclm*-KO, the levels of oxidative stress, as quantified by 8-oxo-2’-deoxyguanosine, increased with age from P20 to P180 ^16^. High levels of 8-oxo-2’-deoxyguanosine appeared to be present in mitochondria and was accompanied with compromised mitophagy ^17^. In *gclm*-KO brains, the upregulation of NAD^+^ (at P20 and P90) and RR (with a peak value at P90) may reflect an adaptation to genetically induced oxidative stress. The sharp decrease of RR in *gclm*-KO mice older than P90 could indicate that such compensatory mechanism via upregulation of NAD^+^ declines in adult *gclm*-KO brains, leading to an exacerbation of the deleterious effects of oxidative stress such as a decreased number of PVIs in the ACC after the age of P90 ^16^. Noteworthy, although PVI maturation appears to be delayed at early postnatal period (P5 - P10) ^16^, it reaches the level of WT mice from peripuberty to young adulthood (P20 - P90) in the ACC, which may partially be attributed to redox homeostasis compensation associated to the NAD^+^ upregulation up to P90 ^27^. Indeed, NAD^+^ has been shown to play a pivotal role in DNA repair and protein deacetylation. High levels of NAD^+^ may promote health and extend lifespan ^28^. The exogenous promotion through supplementation in NAD^+^ precursor-like niacin ^29^, nicotinamide ^30^, nicotinamide mononucleotide ^31^ or nicotinamide riboside ^32^ has been shown to be effective in cell protection by suppressing the oxidative stress in a pathology associated with mitochondrial dysfunction ^33^.

Furthermore, although GABA levels were not altered through development, they are positively correlated with NAD^+^ at P20, a critical time window during which PVIs and brain circuitry undergo intense development and maturation. This suggests that the up-regulation of NAD^+^ may be associated to the maintenance of the GABA level in *gclm*-KO, to prevent deficits in the development of the inhibitory system. Meanwhile, the increase in NAD^+^ was also associated with an increase in Gln at P20. In fact, the last reaction step in the *de novo* synthesis and in some of the NAD^+^ salvage pathways, is the amidation of nicotinic acid adenine dinucleotide, which is achieved by the glutamine dependent NAD^+^ synthetase ^34^.

Taken together, NAD^+^ may serve as a potential preventive treatment target, especially for high risk populations. Here, the assessment of NAD^+^ and RR has been limited to the prefrontal cortex. However, *gclm*-KO displayed oxidative stress and PVI deficits in a temporal and regional specific manner, with PVI anomalies appearing earlier in subcortical regions (i.e. thalamic reticular nucleus, globus pallidus and hippocampus) than in the ACC ^16^. Therefore, a further investigation of the regional and temporal alterations of NAD^+^ and RR and their association with PVI deficits would be extremely valuable to guide preventive time window of NAD^+^ associated treatment.

### Energy metabolism and physiological regulation

During early life, the brain undergoes extensive development and maturation, and high energy is required to form new membranes and myelin sheets around axons. As compared to age-matched WT, the brain of young *gclm*-KO mice (at P20) shows an exacerbated increase in metabolic activity as highlighted by high energy phosphate, ATP (Fig. 3). Intracellular pH, which is tightly associated with cerebral energy status, was also significantly elevated in *gclm*-KO mice at P20, implying reductions in [H^+^] to facilitate ATP synthesis and maintain energy homeostasis as observed in a functional study ^35^. The temporal energy buffer PCr, which supports local energy needs, also increases in *gclm*-KO mice at P20, suggesting that excess ATP is stored to maintain energy homeostasis. In addition, the ratio PME/PDE, reflecting membrane synthesis/degradation, is also significantly higher at P20 in *gclm*-KO mice relative to WT mice, which is mainly driven by the elevated PME (Fig. S2), reflecting the consequence of oxidative damage in membrane lipids that would require more synthesis or repair of membrane elements. This might be also partially responsible for the increase in energetic demand, as ATP is required for membrane turnover.

In patients with SZ, neither ATP nor phosphocreatine (PCr) was consistently affected ^36^. But a magnetization transfer experiment revealed that the reaction rate between PCr and ATP, catalyzed by creatine kinase, was decreased in the ACC of patients ^37^. Here, we established a link between redox state and energy metabolism, however, the enzymatic activity measurement of creatine kinase and ATP synthase would provide additional light on these mechanisms and would constitute a further step in the comprehension of the *gclm*-KO physiology. PCr correlated positively with NAD^+^ only in *gclm*-KO at P20, which suggests a potential adaptation of creatine kinase activity with the upregulation of NAD^+^.

The negative correlation of NADH with Pi^int^ is disrupted in the *gclm*-KO, suggesting aberrant mitochondrial ATP production using ADP + Pi in the electron transport chain. The high ATP and the trend for increased PCr in *gclm*-KO are also observed at P90, indicating that an elevated demand in energy persists in young adult *gclm*-KO, although to a lesser extent than in very young mice. Indeed, *gclm*-KO mice present a lower weight, reduced plasma glucose, insulin and hepatic glycogen levels as compared to WT ^21,38^. These observations are consistent, among other possibilities, with a faster metabolism. Of note, ATP was also suggested to have a protective effect against H2O2 induced oxidative damage ^39^.

The fact that the energy metabolite differences between WT and *gclm*-KO vanish at P40 could be related to a lower energy demand, which is accompanied by a reduction of Gln + Glu at that time of development when synaptic pruning occurs. Therefore, the energy demand may not be as high as in younger and older animals. Our study depicted high Gln and Glu during the early developmental period (at P20) in *gclm*-KO, as previously shown ^40^. The decrease of Gln + Glu observed in our study at P40 could also be partly the consequence of their use as a substitute to power the TCA cycle ^41–43^. Glutamatergic metabolites appear to be elevated in the prodromal and early stages of schizophrenia ^44^, but unchanged or reduced below normal in chronic or medicated patients.

### UDP(G), glycogenesis and extracellular matrix

UDP-glucose is the precursor for glycogenesis. In the brain of *gclm*-KO mouse, the UDPG signal was drastically reduced at P250. Interestingly, adult *gclm*-KO mice displayed low levels of glycogen in their liver ^21^. Likewise, cultured astrocytes from *gclm*-KO mice brain have lower glycogen levels, while its turnover is higher ^38^. This further supports that the regulation of brain energy metabolism is altered together with reduced NAD redox ratio in fully adult *gclm*-KO. Note that oxidative stress challenge induced by tBHQ in human fibroblasts also causes a significant decrease in UDPG ^45^.

Recently, UDPG signal was proposed to result not only from UDP-glucose, but also from other metabolites such as (by order of prevalence) UDP(G): UDP-N-Acetyl-glucosamine (UDP-GlcNAc); UDP-glucose(UDPGlc); UDP-N-Acetyl-galactosamine (UDP-GalNAc) and UDP-galactose(UDP-Gal) ^46^. These molecules are used for the synthesis of the polysaccharide chains of the extracellular matrix that are made of alternating sugar/uronic acid and amino-sugars ^47^. Therefore, the drop of UDPG signal in the brain of adult *gclm*-KO mice might reflect compromised integrity of the perineuronal net, the extracellular matrix enwrapping PVI, as a consequence of chronic oxidative stress. Noteworthy, the extracellular matrix and its homeostasis also appears to be abnormal in patients ^45,48^. The apparent decrease of UDP(G) in full adulthood of *gclm*-KO mice is intriguing, however, further investigation is required, as the spectral SNR achieved in small animal brains is still insufficient to establish which UDP(G) component is altered.

As a limitation in NAD quantification, the UDPG signal overlaps with NAD and has been shown to have an impact on NAD measurements ^49^. Thus, the inclusion of UDPG in the quantification lowers preferentially the NADH content and leads to a higher RR. In the present study, similar results were observed for *gclm*-KO and WT mice on pooled age summed spectra. The NADH values were reduced and RR increased (Table. 1S). The decrease of the RR from P90 to P250 in *glcm*-KO remains nevertheless consistent with the results without inclusion of UDPG and is even more pronounced with the inclusion of UDPG in the analysis.

This study revealed that deficits in redox regulation is associated with a dynamic neurodevelopmental trajectory, leading to specific temporal alterations in NAD redox state and neuroenergetics during the neurodevelopment. The upregulation in NAD^+^, and its highly associated regulations in energy metabolism and in neurotransmission at early developmental period, may be neuroprotective under oxidative stress. This mechanistic insight holds great promise concerning potential preventive treatment targets for at-risk individuals.

## Acknowledgements

We thank Dr. Med. Vet. Stefanita Octavian Mitrea (CIBM) for helpful technical support, Dr. Fulvio Magara of the Centre d’Etudes du Comportement, Center for Psychiatric, CHUV, for support in animal facilities. We also thank the support from Centre d’Imagerie BioMédicale (CIBM) of the UNIL, UNIGE, HUG, CHUV, EPFL, the Leenaards and Jeantet Foundations.

## Author Contributions

L.X. and K.D. designed and directed the study. R.S. and L.X. implemented the method and collected the data. R.S. analyzed data and prepared the figures. R.S. and L.X. wrote the manuscript with the input from K.D. and P.S.

