## Supplementary material for "Developmental changes in cerebral NAD and neuroenergetics of an antioxidant compromised mouse model of schizophrenia": Fig. S1

**Fig. S1: Water content of cerebral cortex in WT and *gclm*-KO mice.**

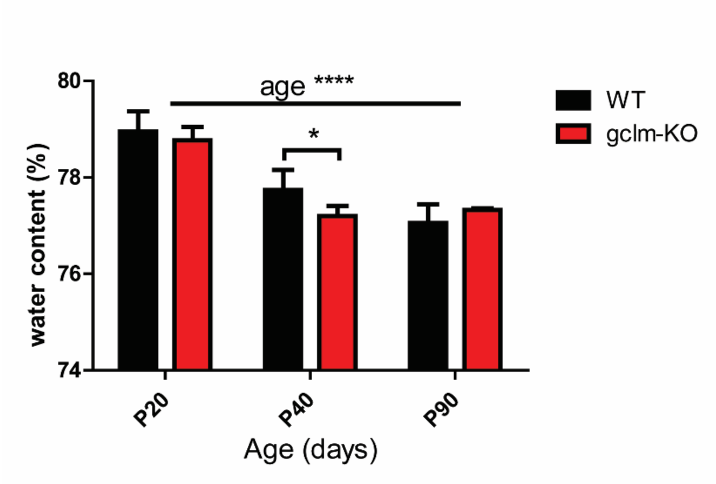


Age dependent cortex water content (%) in WT and *gclm*-KO mice. A significant decrease of water content upon development was observed in both genotypes, where usual $^1$H MRS studies are assuming a constant value. A significant decrease of water content was also seen at P40 in *gclm*-KO mice in comparison to WT.

**Fig. S2: Concentrations of PME and PDE in *gclm*-KO and WT mouse brain during brain development.**

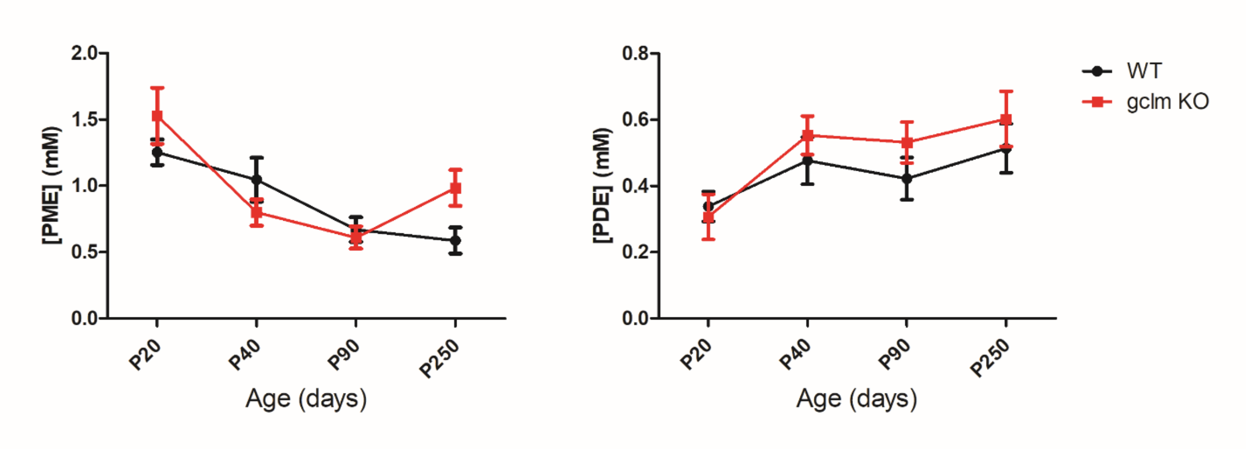

**Table 1S** Levels of UDPG, $NAD^+$, NADH, RR and total NAD obtained with inclusion of UDPG in the quantification from summed spectra at P20, P40, P90 and P250.

| AGE [DAYS] | GENOTYPE | $NAD^+$ [mM] | NADH [mM] | UDPG [mM] | RR [-] | TOTAL NAD [mM] |
|---|---|---|---|---|---|---|
| **P20** | WT | 0.352 | 0.194 | 0.147 | 2.033 | 0.545 |
| | *Gclm*-KO | 0.469 | 0.103 | 0.199 | 5.651 | 0.572 |
| **P40** | WT | 0.368 | 0.155 | 0.185 | 2.422 | 0.523 |
| | *Gclm*-KO | 0.438 | 0.114 | 0.168 | 3.986 | 0.552 |
| **P90** | WT | 0.360 | 0.108 | 0.125 | 3.185 | 0.468 |
| | *Gclm*-KO | 0.622 | 0.021 | 0.167 | 32.745 | 0.644 |
| **P250** | WT | 0.408 | 0.043 | 0.206 | 8.880 | 0.451 |
| | *Gclm*-KO | 0.394 | 0.144 | 0.063 | 3.056 | 0.538 |
